## Supplemental Figures S1 and S2 for "Follicular helper- and peripheral helper-like T cells drive autoimmune disease in human immune system mice"

Figure S1: Ki67 expression in splenic Tph and Tfh cells at 30 weeks post-transplantation. Data are summarized for Mu/Hu (n=3) and Hu/Hu (n=2) mice.

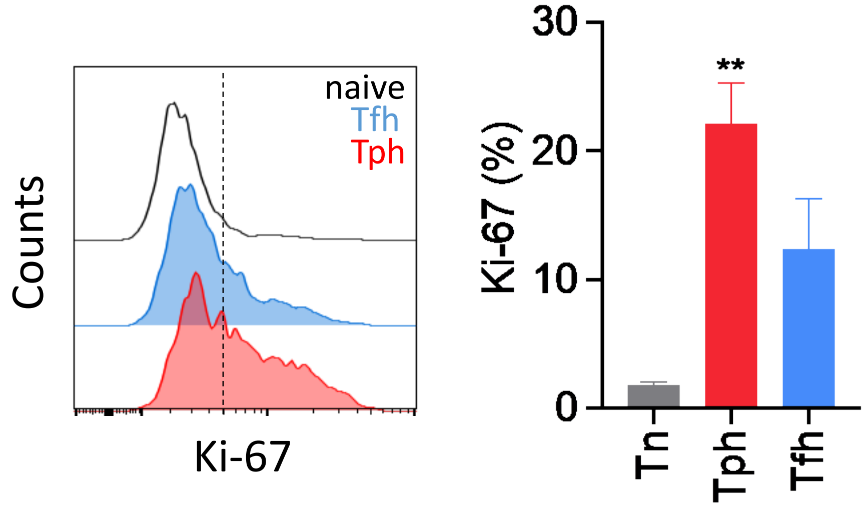

Figure S2: Frequencies of human T and B cell subsets in spleens of adoptive recipients of T cell subsets from Hu/Hu and Mu/Hu mice

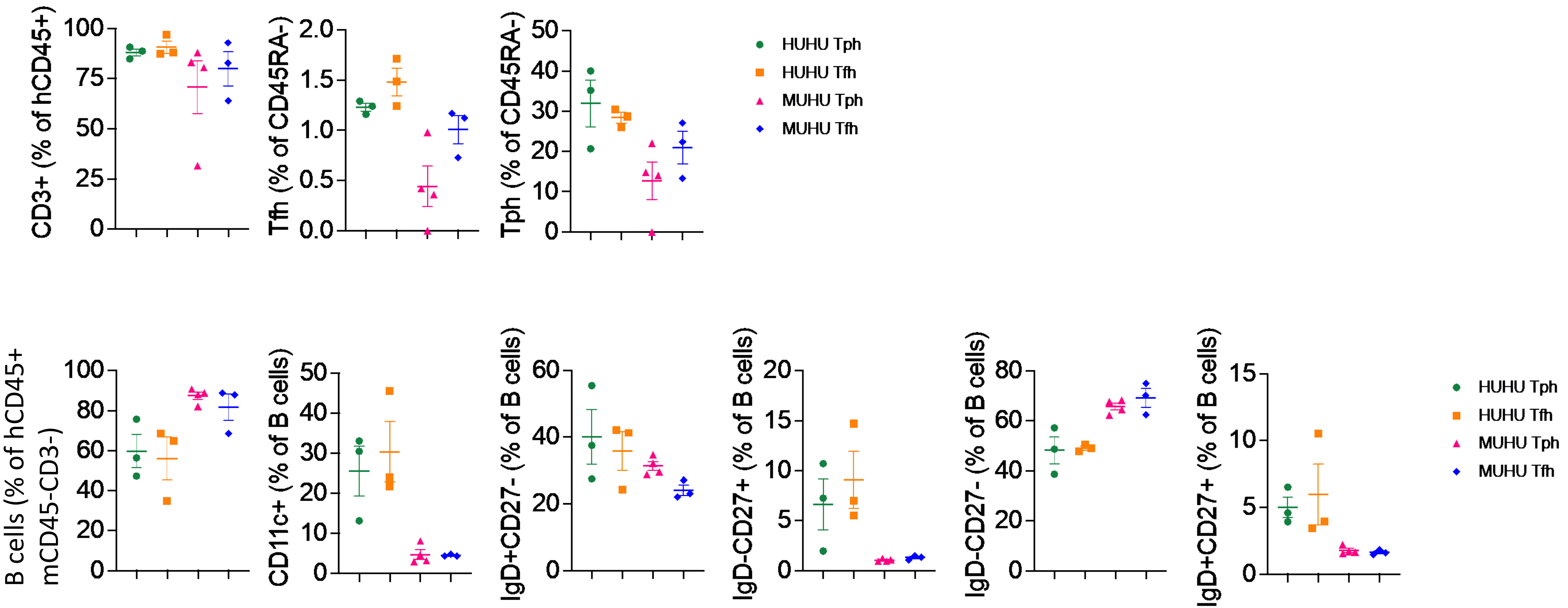
